# Transcription initiation profiling defines the regulatory logic of astrocyte gene regulation

**DOI:** 10.64898/2026.05.03.722406

**Authors:** Ashley Kumar, Yanning Zuo, Numaan Formoli, Daimeng Sun, Narayan Pokhrel, Carlos Guzman, Sven Heinz, Christopher Benner, Francesca Telese

**Affiliations:** Department of Psychiatry, University of California, San Diego Department of Psychiatry, University of California, San Diego; Department of Medicine, University of California, San Diego Department of Psychiatry, University of California, San Diego

## Abstract

Astrocytes are central regulators of neuroinflammation, yet the mechanisms by which they convert common inflammatory signals into cell type-specific transcriptional responses remain poorly understood. Here we mapped transcription initiation genome-wide in primary mouse astrocytes stimulated with interleukin-1B (IL-1B) and defined the active regulatory elements that drive astrocyte reactivity. We find that inducible enhancer transcription in astrocytes is encoded by a transcription-initiation grammar in which lineage-restricted transcription factors, particularly NFIA and TEAD4, cooperate with inflammatory transcription factors such as NF-κB, AP-1 and IRF to drive stimulus-dependent transcription activation. The motifs of these inflammatory transcription factors show a strong positional bias upstream of induced transcription start sites, supporting their direct role in controlling initiation upon stimulation. Moreover, NF-κB and TEAD4 motifs are preferentially associated with sites showing altered patterns of transcription initiation in response to inflammatory stimulus. Comparison with stimulated macrophages revealed that, despite substantial overlap in induced genes, astrocytes exhibit a largely distinct enhancer repertoire, indicating that shared inflammatory signals are interpreted through cell type-specific regulatory landscapes. Finally, transcribed astrocyte regulatory elements are functionally conserved in human astrocytes and are enriched for genetic risk variants associated with neurological disorders. Together, these findings define a cell type-specific regulatory logic for astrocyte inflammatory responses and link astrocyte enhancer regulation to human disease susceptibility.

## Introduction

Cell type-specific gene expression is the fundamental mechanism defining cellular identity and enabling specialized responses to developmental and environmental cues. The regulatory mechanisms governing these programs involve interactions between transcription factors (TFs), chromatin accessibility, and spatial organization of regulatory elements, such as enhancers(1, 2). Critically, the initiation of transcription occurs at transcription start sites (TSSs), where the precise selection of the initiation nucleotide is tightly coupled to the binding of key TFs and their spacing relative to the TSS(3). This positional code represents a fundamental but poorly understood layer of gene regulation, as it determines not only whether transcription occurs, but where and how precisely it initiates. A key functional readout of this process is the production of enhancer RNAs (eRNAs) at active regulatory elements, rapidly degraded noncoding transcripts whose transcription correlates with target gene activity and reflects the engagement of RNA polymerase II (RNAPII) at distal regulatory sites(4–7). While conventional approaches such as ATAC-seq can identify broad regions of open chromatin, they cannot resolve the single-base locations of transcriptional initiation or the positional relationships between TF motifs and TSSs that govern initiation site selection. This leaves a critical mechanistic gap in our understanding of how cell type-specific and stimulus-dependent regulatory codes are encoded in the genome and decoded during gene activation.

This knowledge is particularly critical in the context of neuroinflammation. Astrocytes are the most abundant cells in the brain(8, 9), acting as central regulators of brain homeostasis. Upon activation, a state termed reactive astrogliosis, astrocytes can exert both protective or pathogenic effects depending on context(10–13). Prior transcriptomic and epigenomic studies have revealed widespread changes in regulatory elements during astrocyte activation(13–20); however, the mechanisms by which inflammation-responsive TFs are recruited to astrocyte-specific regulatory elements remains poorly defined. In contrast, macrophages represent a well-characterized system for myeloid-lineage-driven immune response and are the classical model for studying inflammatory gene regulation(21–23).

While astrocytes arise from neuroepithelial progenitor cells and function within the neural tissue(9, 11, 24), macrophages derive from myeloid progenitors and operate in peripheral tissues to recognize pathogens, coordinate tissue repair, and modulate immune responses(25, 26). Both cell types mount robust transcriptional responses to common inflammatory signals, yet their specialized functions demand fundamentally different transcriptional outcomes. Understanding how two functionally distinct cell types decode the same inflammatory signal into cell type-specific gene expression programs therefore represents a powerful opportunity to uncover general principles of stimulus-responsive, lineage-restricted gene regulation.

We addressed this challenge by mapping the regulatory landscapes of primary murine astrocytes at single-base resolution using capped small RNA sequencing (csRNA-seq)(6) in resting and IL-1B-stimulated cells, and comparing them with those previously defined in macrophages(6, 27). The captured nascent short, 5′-capped RNAs produced by active RNAPII at both promoters and promoter-distal regulatory elements, enable genome-wide annotation of astrocyte TSSs within regulatory elements at single-nucleotide resolution and directly measure their transcriptional activity in response to inflammatory stimuli. Together, this allows for high-resolution mapping of TF binding site positioning relative to active TSSs. We characterized the astrocyte-specific regulatory landscape by also mapping binding of lineage-associated TFs and comparing them to the regulatory landscapes of macrophages. Finally, to probe conservation of functionally active regulatory elements in astrocytes and their link to complex traits and diseases, we integrated these genomic data with human data. Together, our results provide a comprehensive characterization of the cell type-specific cis-regulatory grammar in astrocytes, and how it is associated with cis-regulatory genetic variants.

## Results

### Transcriptomic signatures of astrocyte reactivity

To investigate the regulatory mechanisms underlying cell type-specific gene expression in response to proinflammatory stimuli, we first focused on characterizing the transcriptomic signatures of astrocyte reactivity. We isolated primary astrocytes from postnatal day 1 (P1) mouse brains and cultured them in vitro for 21 days to generate mature astrocyte cultures(28) (**Fig. 1A**). These cultures displayed characteristic star-like morphology as confirmed by GFAP immunostaining (**Fig. 1B**). RNA-seq analysis of astrocytes treated with vehicle (n=3) or IL-1B (n=3, 10ng/ml) for 1 hour identified significant transcriptional changes, with 210 genes upregulated and 28 downregulated (Fold Change > 2, FDR < 0.05; **Fig. 1C,D, Supp Table 1, 2**). The top upregulated genes included established proinflammatory mediators such as chemokines and cytokines (e.g., *Cxcl1, Ccl20, Csf2, Il6*, **Fig. S1**)(25). Pathway enrichment analysis(29) confirmed that the differentially expressed genes (DEGs) were primarily associated with TNF signaling, inflammatory responses, and interleukin-1 pathways **(Fig. 1E**). Further validation by RT-qPCR (**Fig. 1F**) reinforced the reliability of these observations. These findings validated the experimental model as reflective of astrocyte-specific reactivity.

**Figure 1:**
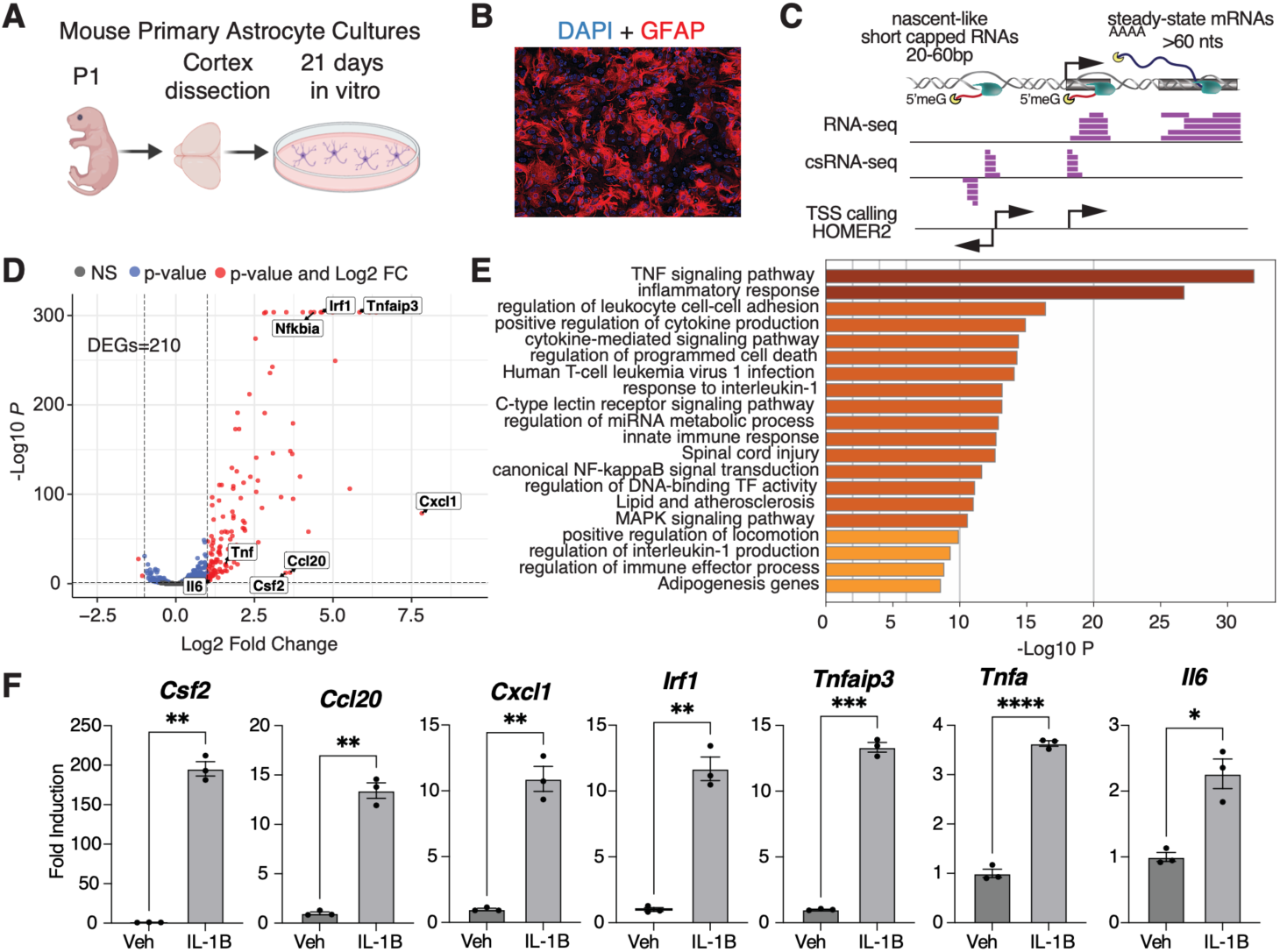
Transcriptomic signatures of astrocyte activity. (A) Schematic of the primary mouse astrocyte cultures. (B) Representative immunofluorescence images of cultured astrocytes stained for GFAP, showing characteristic astrocyte morphology. (C) Overview of RNA-seq and capped small (cs)RNA-seq data generation from astrocytes. The schematic shows the typical distribution of csRNA-seq and RNA-seq at various genomic locations, which allows the identification of Transcriptional Start Site (TSSs) using HOMER2. (D) Volcano plot of RNA-seq differential expression in astrocytes treated with vehicle (Veh) or IL-1B (10 ng/mL, 1 h). (E) Pathway enrichment analysis of IL-1B-induced differentially expressed genes. (F) RT-qPCR validation of selected IL-1B-responsive genes in astrocytes

### Mapping transcribed regulatory elements in astrocytes

To define the active regulatory landscape of astrocytes, we performed csRNA-seq in resting and IL-1B-stimulated cells, collectively identifying 69,435 genomic Transcriptional Start Regions (TSRs; **Supp. Table 1, Fig. S2A**), which represent strand-specific clusters of individual TSSs that mark active regulatory elements. Approximately 38% of TSRs overlapped promoter-proximal sites associated with annotated genes(30) (**Fig. S2B,C**), while the remaining 62% were promoter-distal elements, consistent with widespread enhancer (eRNA) transcription (**Fig. 2A, S1C**). Most TSRs marked regions that were bidirectionally transcribed (82%), and 70% did not produce stable transcripts detectable by conventional RNA-seq data (**Supp. Table 1**), underscoring the sensitivity of csRNA-seq for capturing nascent transcription at regulatory elements.

**Figure 2:**
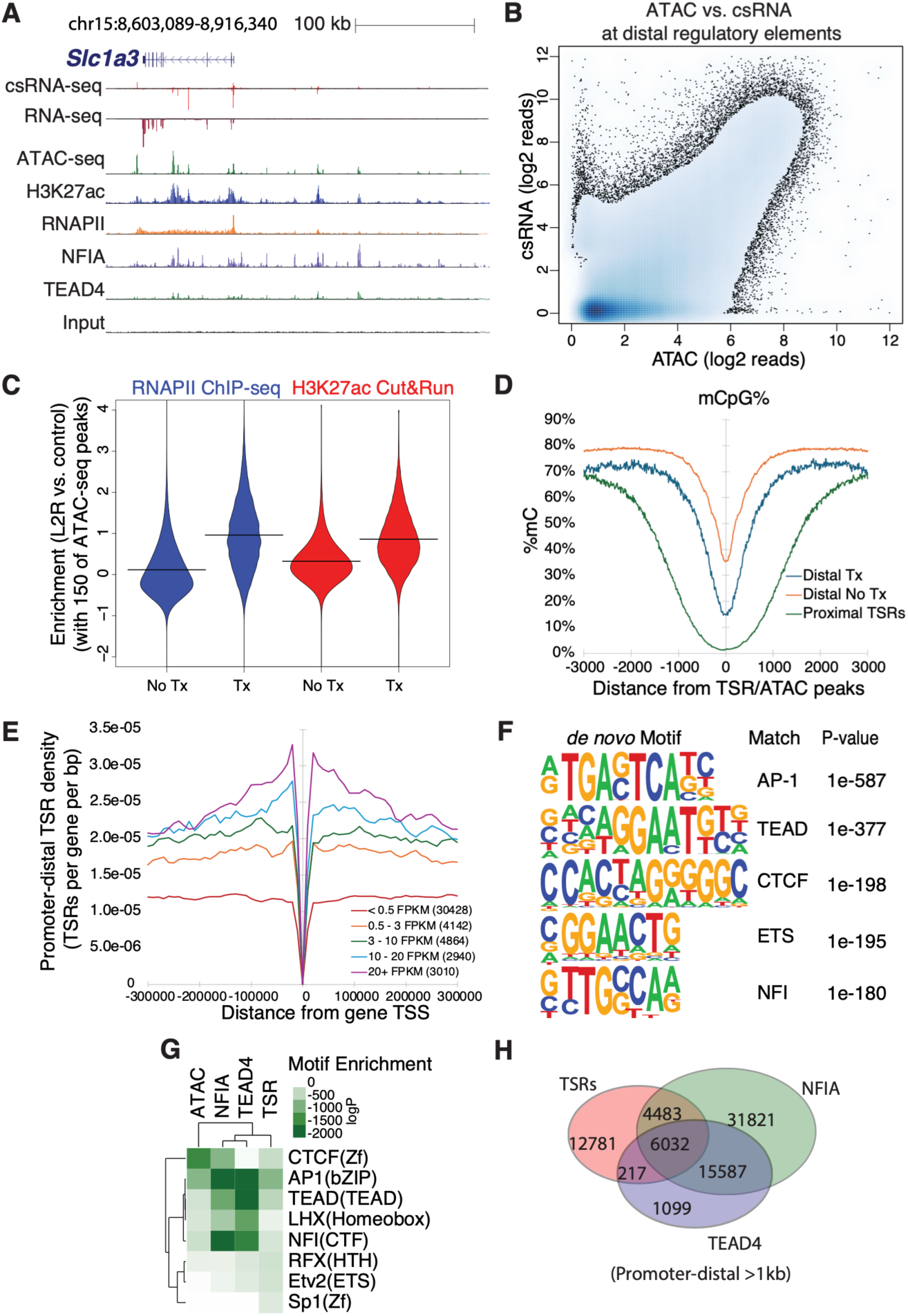
Mapping transcribed regulatory elements in astrocytes. (A) Genome browser view of regulatory regions at a representative astrocyte locus (*Slc1a3*), illustrating correspondence between transcription initiation, open chromatin, active chromatin marks, and TF binding at transcribed regulatory elements. (B) Density scatter plot comparing ATAC-seq and csRNA-seq signal at promoter-distal regulatory elements. (C) Violin plots showing enrichment of RNAPII ChIP-seq and H3K27ac CUT&RUN signal at transcribed (Tx) versus non-transcribed (No Tx) distal ATAC-seq peaks. (D) Average mCpG profiles centered on active regulatory elements defined by TSRs (Transcription Start site Regions) and ATAC peaks with (Tx) and without (No Tx) evidence of transcription. (E) Average density of promoter-distal TSRs around genes stratified by expression level. (F) Top 5 de novo motifs identified by HOMER in promoter-distal TSRs in resting astrocytes (−150, +50 relative to the primary TSS). (G) Comparison of TF motif enrichment across ATAC-seq peaks, csRNA-seq TSRs, and NFIA and TEAD4 ChIP-seq peaks using HOMER. (H) Overlap of promoter-distal TSRs (>1 kb from annotated TSS) with NFIA and TEAD4 ChIP-seq peaks in resting astrocytes.

Integration of the csRNA-seq data with ATAC-seq and H3K27ac ChIP-seq datasets from astrocytes(20) revealed that most TSRs overlapped with open chromatin (83%) and the active enhancer mark H3K27ac (87%) (**Supp. Table 1**), establishing them as bona fide active regulatory elements (**Fig. 2A, Fig. S2D**). While chromatin accessibility and transcription initiation were generally correlated at distal regulatory elements, a substantial fraction of accessible regions lacked detectable transcription (17%, **Fig. 2B, S2E**), demonstrating that accessibility alone is insufficient to distinguish poised from active regulatory elements. Notably, transcribed distal ATAC-seq peaks (>3 kb from annotated TSS) exhibited significantly higher enrichment for RNAPII and H3K27ac relative to non-transcribed accessible peaks (**Fig. 2C**), confirming that active enhancer transcription is coupled with H3K27ac and RNAPII engagement(7, 31–33). Consistent with this, analysis of whole-genome bisulfite sequencing(34) revealed pronounced depletion of methylation of CG dinucleotides (5mCpG) at TSRs relative to non-transcribed accessible regions (**Fig. 2D**). The depletion of 5mCpG was strongest at promoter-proximal TSRs, consistent with extensive demethylation at actively transcribed promoters and relatively intermediate methylation at distal enhancer elements(35–37) (**Fig. 2D**). Depletion of non-CpG-cytosine methylation (mCH), an epigenetic modification typically enriched in neurons, was also observed at astrocyte TSRs genome-wide (**Fig. S2F**). Furthermore, highly expressed genes were flanked by a greater number of promoter-distal TSRs than lowly expressed or silent genes (**Fig. 2E**), reinforcing the link between enhancer transcription and target gene activation.

We next asked which TFs govern enhancer selection and activity in astrocytes. *De novo* motif enrichment analysis of promoter-distal TSRs using HOMER(3, 38) revealed strong enrichment for AP1, TEAD, CTCF, ETS, and NFI TF families (**Fig. 2F**), many of which have established roles in astrocyte identity and reactivity(39–41). Notably, TEAD and NFI motifs ranked among the top enriched motifs, providing rationale for experimental validation by ChIP-seq. While NFIA is a known astrocyte transcriptional regulator(42–44), the role of TEAD4 in astrocytes has remained unclear despite its established function in neural progenitor cells(45). ChIP-seq identified 68,720 NFIA and 25,104 TEAD4 binding sites in untreated astrocytes (**Fig. S2G,H**), with motif analysis confirming high enrichment for their canonical motifs (**Fig. 2G**). While each factor occupied both transcribed and non-transcribed accessible sites individually, sites co-bound by both NFIA and TEAD4 were more than twice as likely to be transcribed (28% vs. 12.5%, **Fig. 2H, S2G**), suggesting their co-binding may contribute to transcriptional activation of regulatory elements.

In summary, these findings distinguished between poised and active enhancers in astrocytes and identified TFs that contribute to enhancer activation in astrocytes.

### Differential regulation of regulatory elements by interleukin-1B

To investigate the regulatory code underlying the inflammatory response in astrocytes, we analyzed how the transcription initiation landscape changes following IL-1B stimulation. We identified 4101 upregulated (Log2FC, >1; FDR, <0.05) and 1845 downregulated (Log2FC, <-1; FDR, <0.05) TSRs compared to vehicle-treated controls (**Fig. 3A**). csRNA-seq captured approximately 28-fold more differentially regulated features than RNA-seq (5,946 TSRs vs. 210 genes, **Fig. 3A vs. 1C**), underscoring its enhanced sensitivity. The upregulated TSRs were enriched near genes involved in IL-1B signaling and astrocyte reactivity, such as *Ccl2* (**Fig 3B**) and *Cxcl10* (**S3A**)(46). Functional annotation using GREAT(47) confirmed their association with immune regulation, cytokine production, and inflammatory response pathways (**Fig 3C**). Notably, 56% (2,293 out of 4,101) of upregulated TSRs mapped promoter-distally (>3 kb from annotated promoters) and co-clustered with other induced TSRs into spatial domains of approximately 50 kb (**Fig. 3D**), consistent with coordinate enhancer activation across extended genomic regions during the inflammatory response(48–50).

**Figure 3:**
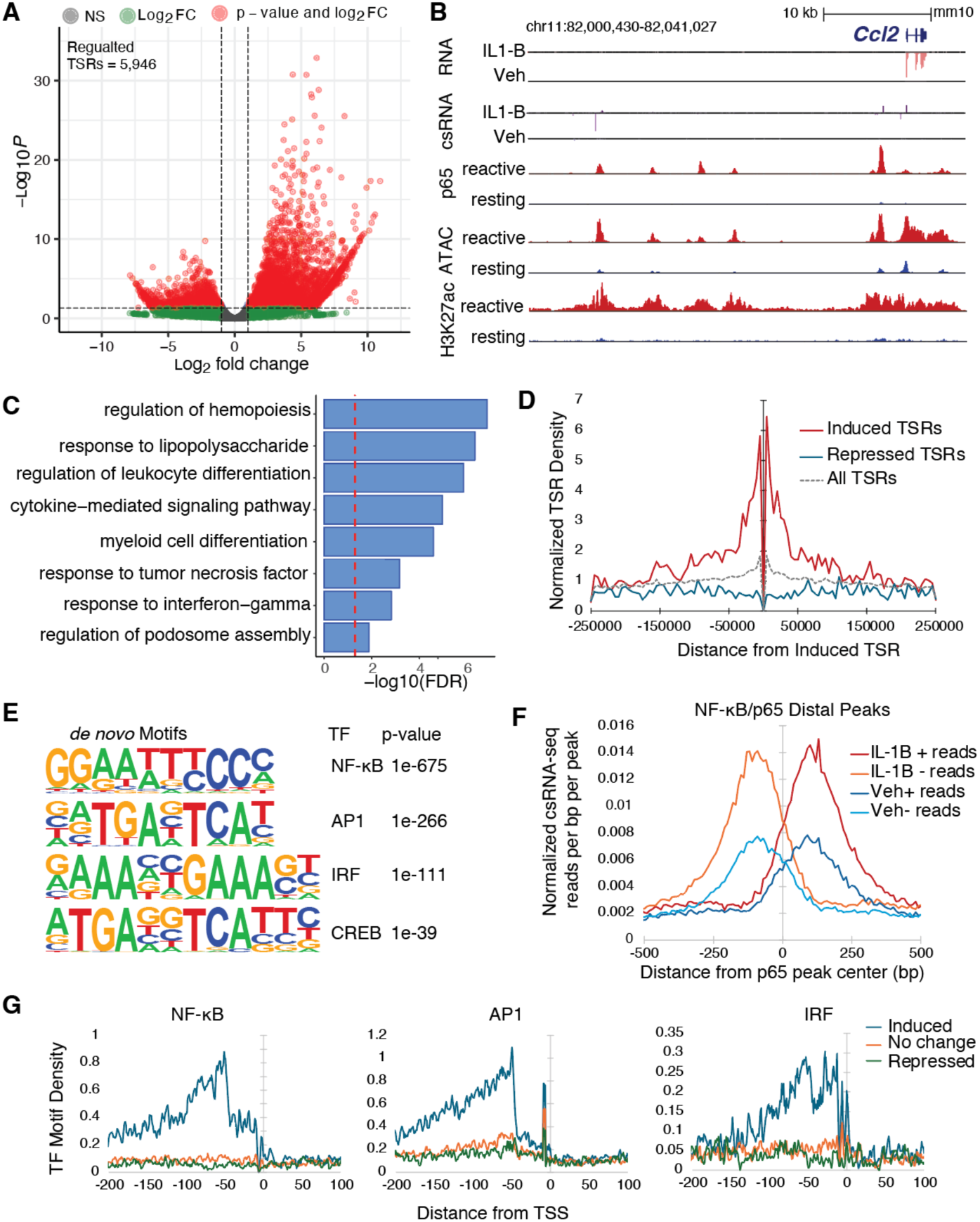
Differential regulation of regulatory elements by interleukin-1B. (A) Volcano plot of differentially regulated csRNA-seq signal at astrocyte TSRs following IL-1B stimulation, identifying induced and repressed regulatory elements. (B) Genome browser tracks depicting induction of promoter and distal regulatory enhancer transcription by csRNA-seq at the *Ccl2* locus. (C) GREAT functional enrichment analysis of IL-1B induced TSRs. (D) Spatial clustering of IL-1B induced TSRs, depicting the density of TSRs in each category adjacent to Induced TSRs. (E) De novo motif enrichment analysis of IL-1B induced TSRs by HOMER. (F) Average csRNA-seq signal centered on promoter-distal NF-κB/p65 peaks, showing increased bidirectional transcription at p65-bound distal elements after IL-1B stimulation, consistent with eRNA induction. (G) Spatial density of NF-κB, AP1, and IRF motifs relative to TSRs (TF binding sites per bp per TSS), showing upstream enrichment of these motifs at IL-1B-induced TSRs compared with unchanged or repressed TSRs.

Comparison of csRNA-seq and RNA-seq profiles revealed that IL-1B regulates transcription through distinct mechanisms (**Fig. S3B**). While many genes showed concurrent induction of both promoter initiation and mRNA levels, others fell into two additional classes. A subset of genes showed strong induction at the mRNA level with minimal changes in initiation frequency (**Fig. S3B**, along the y-axis), characteristic of genes regulated primarily at the level of transcriptional elongation rather than initiation and consistent with the release of paused RNAPII at promoters of pre-loaded immediate response genes such as *Junb* (**Fig. S3C,D**)(51–53). Conversely, a large number of genes showed strong increase in transcription initiation with minimal corresponding change in mRNA levels (**Fig. S3B,** along the x-axis), suggesting that, for these genes, initiation is necessary but not sufficient for productive elongation, and additional regulatory steps are necessary for RNAPII to transition from initiation to productive elongation at these sites.

To identify the TFs driving the initiation response, *de novo* motif enrichment of IL-1B-induced TSRs revealed strong enrichment for NF-κB, AP1, and IRF TF families (**Fig. 3E**), all known mediators of neuroinflammatory pathways(18). Validation using previously identified NF-κB/p65 binding sites in reactive astrocytes(20) confirmed that 33% of IL-1B-regulated TSRs coincided with p65 binding (**Supp. Table 3**). Promoter-distal p65 binding sites exhibited elevated eRNA levels following stimulation, supporting a direct role for NF-κB in driving transcription initiation during the inflammatory response (**Fig. 3F**). Leveraging the single-base resolution of csRNA-seq, we next examined the spatial organization of enriched TF motifs relative to TSSs. NF-κB, AP1, and IRF motifs were preferentially positioned upstream of IL-1B-regulated TSSs compared to uninduced or repressed TSSs, with peak enrichment at approximately −50 bp relative to the TSS that decayed with increasing distance (**Fig. 3G**). This upstream positional bias was consistent across all three TF families, supporting their direct mechanistic role in RNAPII recruitment and TSS selection during neuroinflammation.

Beyond changes in overall transcriptional activity, changes in initiation site usage may indicate the repositioning of RNAPII as a result of altered TF activity or co-regulatory complex engagement with regulatory DNA. We next asked whether IL-1B stimulation alters the distribution of initiation events within TSRs in a stimulus-dependent manner, independent of changes in total transcriptional output. Using WIP scoring(54) we identified 1565 TSRs exhibiting significantly altered initiation patterns without corresponding changes in overall transcription output (p < 0.05, **Fig. S3E,F, Supp Table 1**). Motif enrichment at these sites revealed a potential mechanistic divergence among TFs (**Fig. S3G**). NF-κB motifs were enriched at TSRs with both altered overall activity and TSS patterns, consistent with its rapid nuclear translocation upon activation and direct role in establishing new initiation sites(55, 56). TEAD motifs were most selectively enriched at sites with altered TSS usage patterns upon IL-1B stimulation, suggesting a role in remodeling initiation architecture during inflammation. In contrast, NFI motifs were preferentially enriched in sites with changes in overall activity rather than initiation architecture, suggesting that NFI acts primarily to modulate transcriptional outcome rather than TSS selection. These findings prompted us to examine the binding profiles of NFIA and TEAD4 following IL-1B stimulation. ChIP-seq analysis showed a stimulus-dependent redistribution of both factors, identifying 22,872 new NFIA and 48,558 new TEAD4 binding sites following stimulation, with TEAD4’s large increase in binding consistent with its potential influence on initiation patterns (**Fig S3H**). Newly bound sites were strongly enriched for co-binding with NF-κB, and motif analysis confirmed enrichment for NF-κB, AP1, and IRF motifs at these sites (**Fig. S3I**) indicating that lineage TFs are actively redistributed to inflammatory response elements upon stimulation.

Collectively, these findings emphasize how the changes in TSS positions and levels reveal evidence that key TF motifs, including stimulus-dependent (NF-κB, AP1, and IRF) and lineage-defining (NFIA, TEAD4) TFs, play crucial roles in activating the astrocytic regulatory program during neuroinflammation.

### Cell-type-specific enhancer landscapes underlie divergent inflammatory gene programs

To gain insights in the regulatory logic underlying cell type-specific inflammatory responses, we compared astrocytes to macrophages. Each cell type derives from distinct developmental progenitors, but both serve as key innate immune responders within their respective tissues. Supporting this, IL-1B-stimulated astrocytes exhibited substantial overlap (∼50%) in upregulated genes with KLA-stimulated mouse bone marrow-derived macrophages (BMDMs), as assessed by matched RNA-seq and csRNA-seq(6, 27) (**Fig 4A, S4A, Supp Table 2, 3**). Consistent with this functional convergence, both cell types showed enrichment for the same stimulus-dependent TF motifs at induced TSRs, including NF-κB, AP1, and IRF family motifs(6). Despite the large overlap in induced genes and enriched TF motifs, comparison of induced TSRs revealed largely divergent regulatory landscapes (**Fig. 4B**, **Fig. S4B**). Only 25% of significantly upregulated TSRs overlapped between astrocytes and BMDMs. This is also reflected at gene loci induced in both systems. Of the 580 stimulus-induced TSRs located within 25 kb from the promoters of common response genes (101 genes total), only 19% were induced in both cell types, while 17% were astrocyte-specific and 63% were BMDM-specific. For example, *Tnfaip3* is induced by both cell types as measured by RNA-seq and csRNA-seq at the promoter. However, promoter-distal induced TSRs found upstream of the promoter are found at different locations in the two cell types despite that both sets recruit NF-κB p65 (**Fig. 4C**).

**Figure 4:**
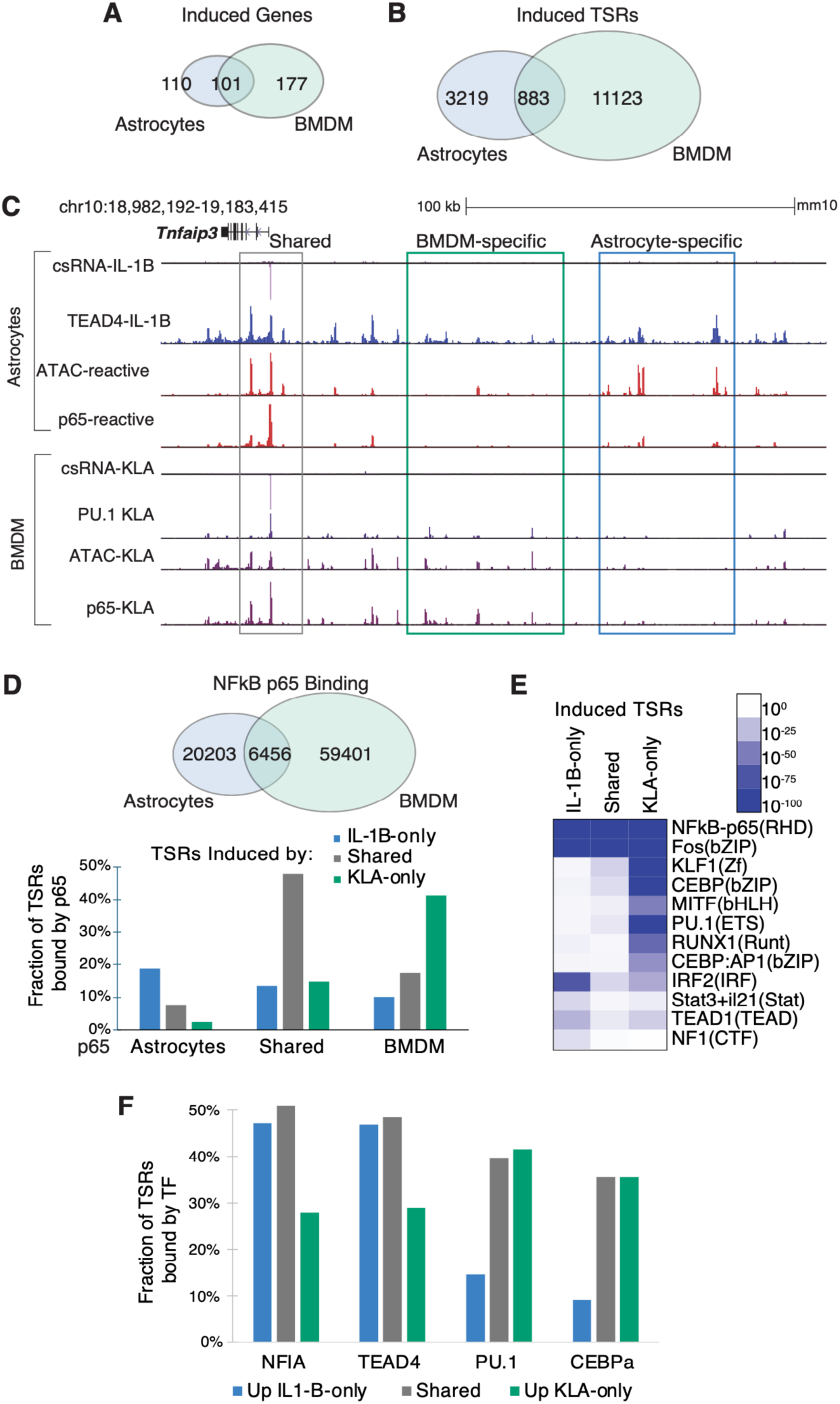
Cell type-specific enhancer landscapes underlie divergent inflammatory gene programs. (A) Venn diagram showing overlap between IL-1B-induced genes in astrocytes and KLA-induced genes in bone marrow-derived macrophages (BMDMs). (B) Venn diagram showing overlap between induced TSRs in astrocytes and BMDMs, revealing largely distinct stimulus-responsive enhancer landscapes despite partial overlap in induced genes. (C) Genome browser view of *Tnfaip3* locus illustrating that astrocytes and BMDMs induce the same gene but use different enhancers upstream a shared promoter. (D) Top: Venn diagram showing overlap of NF-κB binding sites between activated astrocytes and BMDMs. Bottom: Fraction of astrocyte-specific, shared, and BMDM-specific p65 peaks associated with TSRs induced in astrocytes only, in both cell types, and in BMDM only. (E) Heatmap of de novo motif enrichment at cell type-specific induced TSRs, showing enrichment of lineage-associated motifs for each cell type. (F) Fraction of induced TSR classes bound by lineage-associated TFs, showing preferential association of astrocyte-induced TSRs with NFI and TEAD4 and macrophage-induced TSRs with PU.1 and CEBPɑ.

To understand the basis of this regulatory divergence, we compared NF-κB p65 ChIP-seq data from activated astrocytes(20) and BMDMs(27). This analysis revealed largely non-overlapping binding cistromes, consistent with the divergent TSR repertoires described above (**Fig. 4D**). Cell type-specific induction of eRNAs at these sites further supported the direct role of NF-κB in driving transcription initiation in a cell type-restricted manner (**Fig. S4A**). Moreover, NF-κB binding was preferentially observed at open chromatin regions that were established in a cell type-specific manner prior to stimulation (**Fig. S4B**), indicating that the pre-existing chromatin landscape substantially contributes to where NF-κB is recruited. DNA motif analysis of cell type-specific response TSRs revealed enrichment for astrocyte-associated TFs (TEAD, RFX, NFI) at astrocyte-specific sites and BMDM-associated TFs (PU.1, CEBP) at macrophage-specific sites, directly implicating lineage factors in establishing the cell type-specific enhancer repertoire available for inflammatory activation (**Fig. 4E**). Consistent with this model, ChIP-seq analysis demonstrated that NFIA and TEAD4 cistromes in IL-1B stimulated astrocytes, and PU.1 and C/EBPβ binding in KLA-stimulated BMDMs, overlapped with cell type-restrictedly induced TSRs (**Fig. 4F**). Furthermore, NF-κB/p65 binding was often accompanied by increased occupancy of lineage TFs following inflammatory activation (**Fig. S4C**). For example, TEAD4 occupancy increases at NF-κB binding sites following IL-1B treatment, suggesting lineage TFs and inflammatory TFs are co-recruited in a cooperative manner that reinforces cell type-specific enhancer activation, consistent with previous observations in macrophages and B cells(27, 38).

These findings support a model in which cell type-specific gene regulatory networks emerge from cooperation between lineage-determining and stimulus responsive TFs.

### Conservation of astrocyte regulatory elements and enrichment for disease risk variants

To establish the translational relevance of our mouse astrocyte regulatory map, we first tested whether transcribed regulatory elements identified in mouse astrocytes are conserved and functional in human astrocytes before assessing their enrichment for human disease-associated variants. We therefore compared mouse astrocyte TSRs with publically available epigenomic and transcriptional datasets from human astrocytes. Of the 69,435 TSRs identified in mouse astrocytes, 59,653 mapped to homologous DNA in the human genome (see Methods). 71.3% and 69.0% of mouse TSRs overlapped with open chromatin in human astrocytes, as measured by scATAC-seq from postmortem brains(57) or by ATAC-seq in cultured fetal primary human astrocytes(58), respectively. These rates declined to 42 and 43%, respectively, when restricted to gene-distal, intergenic regions, which is consistent with the accelerated turnover of distal regulatory elements across species(59).

Analysis of nascent RNA-seq data (TT-seq)(58) from human astrocytes revealed strong evidence for eRNA expression from conserved mouse TSRs overlapping human astrocyte ATAC-seq peaks compared to those mapping to inaccessible regions, with a clear enrichment of both sense and antisense nascent transcription centered at the conserved TSR position (**Fig. 5A, 5B, S5A**). This suggests that a subset of mouse astrocyte regulatory elements retains functional transcriptional activity in human cells. Sites active in both mouse and human had higher rates of sequence conservation(60) **(Fig. 5C**) and analysis of AP1, NFI, and TEAD4 motif positioning in human genomic DNA revealed preferential enrichment at conserved, accessible TSRs sites relative to non-accessible sites. In contrast, motifs for CEBP factors, which are not active in astrocytes, were not enriched (**Fig 5D, S5B**), indicating that sequence-level preservation of TF binding architecture contributes to functional conservation across species.

**Figure 5:**
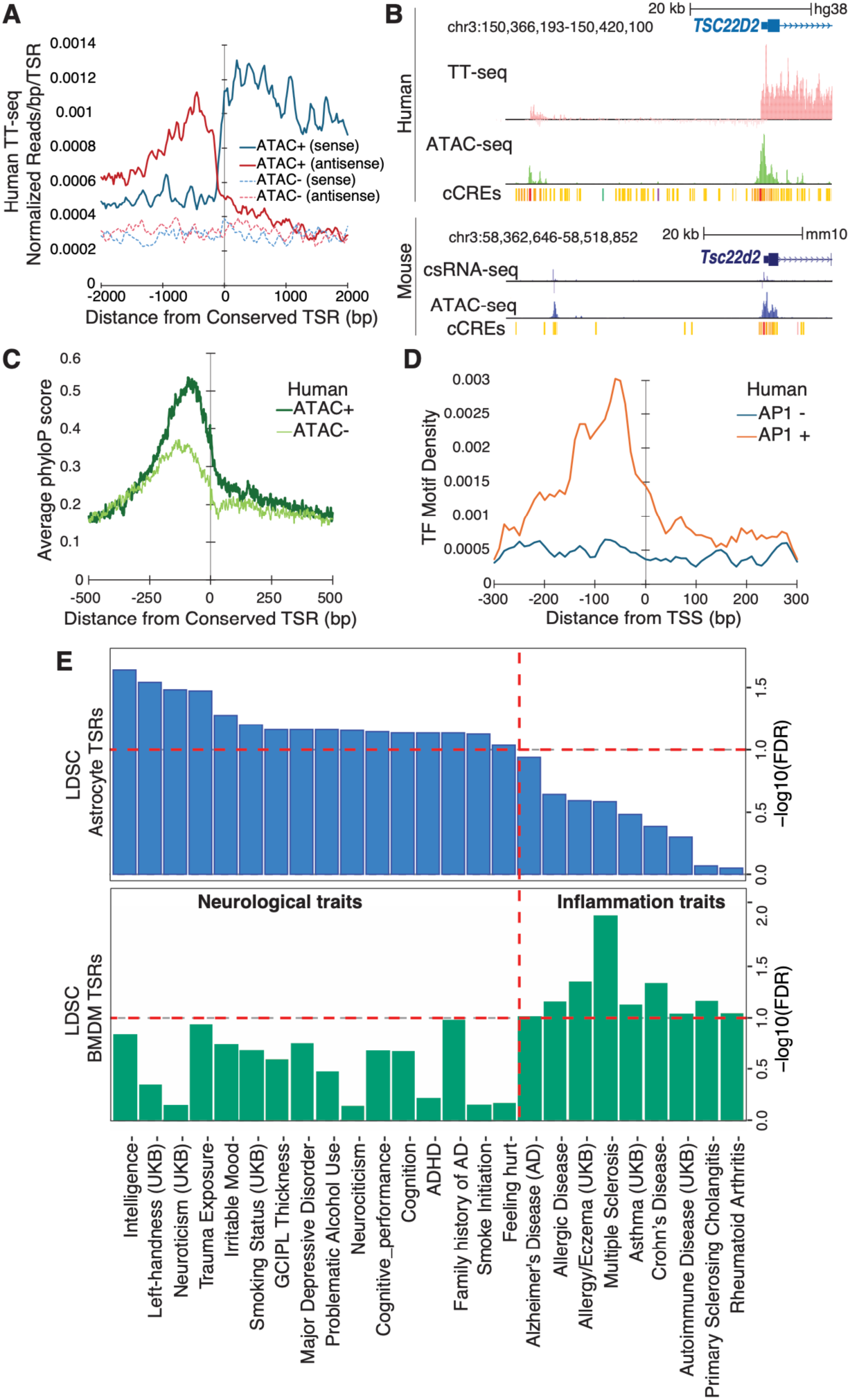
Conservation of astrocyte regulatory elements and enrichment for disease risk variants. (A) Average human astrocyte nascent RNA levels (TT-seq) signal centered on conserved mouse TSRs, separated by those that overlap ATAC-seq peaks in human astrocytes (ATAC+) versus those that do not (ATAC-). (B) Genome browser view of a conserved human and mouse eRNAs upstream of the *TSC22D2/Tsc22d2* gene, showing aligned ATAC-seq, TT-seq/csRNA-seq, and ENCODE candidate cis-regulatory element tracks in human and mouse astrocytes. (C) Genome browser view of a conserved human and mouse eRNAs upstream of the TSC22D2/Tsc22d2 gene, showing aligned ATAC-seq, TT-seq/csRNA-seq, and ENCODE candidate cis-regulatory element tracks in human and mouse astrocytes. (D) Distribution of AP1 motifs density (TF motifs/bp/TSR) around conserved mouse TSR positions in the human genome, comparing sites that are accessible or inaccessible in human astrocytes. (E) LDSC-based enrichment of GWAS signal within conserved mouse astrocyte or macrophage TSRs, highlighting neurological- and inflammation-related trait associations that are preferentially enriched in each cell type.

To further validate the functional relevance of conserved TSRs, we integrated our data with results from a CRISPRi Perturb-seq screen of candidate regulatory elements in human astrocytes(58). Of the 957 regions targeted in their screen, 256 (26.7%) overlapped with conserved mouse TSRs. Further analysis showed that 23.8% of target regions overlapping a mouse TSR were functional enhancers (61/256) compared to 12.0% of non-overlapping regions (84/701), indicating that conserved human sites that are transcribed in mouse were approximately twice as likely to have functional consequences for gene expression than other sites.

Astrocyte-transcribed regulatory elements were next examined for their relevance to human disease genetics. As an initial broad test, we performed a variant-TSR overlap enrichment analysis(61) using 2,025 well-powered GWAS traits from the NHGRI-EBI GWAS catalog, each with more than 20,000 cases(62), and found that 96 neurological traits were significantly enriched in astrocyte TSRs (FDR<5%, **Supp Table 4**). This suggested that astrocyte regulatory elements defined in mouse primary cultures capture conserved genomic features relevant to human CNS disease risk, motivating a more quantitative assessment of trait heritability enrichment.

Next, we applied the more rigorous linkage disequilibrium score (LDSC) regression(63, 64) to quantify the enrichment of GWAS variants from 302 different traits within conserved TSRs identified in mouse astrocytes, using conserved mouse BMDMs TSRs as a reference to control for cell type specificity. We identified 143 significant trait-TSRs associations (FDR < 0.1, **Supp Table 5**, **Fig. 5E, Fig. S6**) of which 65 were enriched in astrocytes and 78 in BMDMs TSRs, highlighting strong cell type specificity in genetic risk enrichment. Astrocyte TSRs were selectively enriched for risk variants linked to brain-related traits and disorders (**Fig. 5E**), including Alzheimer’s disease, trauma exposure, addiction, and bipolar disorder(65–68), consistent with emerging evidence that astrocytes contribute to the pathophysiology of diverse CNS conditions. In contrast, BMDMs TSRs showed strongest enrichment for immune and hematological traits, consistent with their established roles in immunological functions (**Fig. 5E, Fig. S6A**).

Together, these findings demonstrate that transcribed astrocyte regulatory regions identified in mouse primary cultures are conserved in human cells and enriched for genetic risk associated with CNS disorders.

## Discussion

Using csRNA-seq to map transcriptional initiation at single-base-pair resolution in astrocytes, we define transcribed regulatory elements and establish a TSS-based regulatory grammar in which key astrocyte TF motifs influence both transcription initiation efficiency and TSS selection. We demonstrate how lineage-determining TFs direct NF-κB genomic targeting to define cell type-specific inflammatory enhancer repertoires and establish that astrocyte-specific TSRs are selectively enriched for CNS disorder risk variants. Together, our findings reveal that the regulatory logic governing astrocyte-specific inflammatory responses is encoded at the level of transcription initiation itself.

A key result of this study is a spatial regulatory grammar at active TSSs in astrocytes, consistent with recent work demonstrating position-dependent TF function in human regulatory elements(3, 69). Our findings extend this model to the astrocyte inflammatory response. First, NF-κB, AP1, and IRF were preferentially positioned upstream of IL-1B-induced TSSs, indicating that the spatial arrangement of sequence-specific TFs is critical for efficient recruitment of RNAPII and initiation of transcription during neuroinflammation. Second, the internal architecture of TSS usage patterns is itself regulated in a TF-specific manner. Specifically, TEAD and NF-κB motifs were preferentially associated with stimulus-dependent shifts in TSS selection, supporting a model in which certain TFs determine transcriptional precision, directing where within a regulatory element the pre-initiation complex assembles(70–74). In contrast, NFI motifs were primarily associated with changes in overall transcriptional magnitude without corresponding shifts in TSS patterns, consistent with models in which distinct classes of TFs independently regulate RNAPII recruitment and productive elongation(75–77).

The comparison between astrocytes and macrophages revealed that despite substantial overlap in stimulus-induced mRNA transcripts, the underlying enhancer repertoires were predominantly divergent with only ∼25% of induced TSRs shared. This finding supports a mechanistic model by which universal inflammatory TFs such as NF-κB can mediate specialized regulatory programs in different cell types(78). We show that lineage-determining factors establish cell type-specific chromatin landscapes that define the sites to which stimulus-responsive TFs are preferentially recruited. In astrocytes, NF-κB is recruited to sites co-bound by NFI and TEAD, whereas in BMDMs it is directed by PU.1 and C/EBP factors. This mechanism aligns with previous work demonstrating the combination of lineage-specific TFs prime cell type-specific landscapes in macrophages and B cells(38). Our ChIP-seq data showing stimulus-induced co-recruitment of NFIA and TED4 to NF-κB-enriched sites further corroborate this collaborative binding model.

These findings also identify TEAD4 as a previously uncharacterized regulator of astrocyte inflammatory gene expression. TEAD factors canonically function as transcriptional effectors of YAP/TAZ downstream of the Hippo pathway(79–81), and while YAP has been implicated in astrocyte differentiation, it generally attenuates rather than promotes neuroinflammation(81–86). Our findings raise the possibility that TEAD4 contributes to astrocyte inflammatory gene regulation through a mechanism that is at least partially independent of canonical YAP co-activator activity. One candidate mechanism is the TRAF6-YAP-TEAD4 signaling axis, previously described in other cellular contexts(87), which could provide a route for IL-1B to activate TEAD4 in astrocytes independently of Hippo pathway status and warrants further investigations.

Integrating our astrocyte TSR map with human GWAS data reveals that the regulatory code defined here is directly relevant to human disease genetics. Transcribed mouse astrocyte TSRs were approximately twice as likely to correspond to functionally validated enhancers in human CRISPRi screens(15) compared to non-transcribed accessible regions, establishing active transcription as a robust predictor of regulatory function across species. Leveraging this conservation, we show that astrocyte TSRs are significantly enriched for heritability of CNS disorders, including AD, bipolar disorders and neuroticism, while macrophage TSRs were enriched for variants associated with hematological and immune traits, a pattern that directly mirrors the known biological roles of each cell type. The specificity of these associations support the broader principle that disease-associated non-coding variants preferentially disrupt cell type-specific regulatory elements(88–91).

Several limitations of this study should be noted. The regulatory mechanisms identified here are based on cooccupancy and motif evidence, and direct perturbation experiments will be needed to establish causal roles for NFIA and TEAD4 in governing the astrocyte inflammatory response. Additionally, the csRNA-seq experiments were performed in bulk primary astrocyte cultures stimulated with a single cytokine (IL-1B), which may not fully recapitulate the diversity of astrocyte states and regulatory responses present in vivo, where astrocytes are exposed to a combination of signals from neurons, microglia and other cell types, and where epigenomic states may differ from those of cultured cells. Future studies incorporating single-cell transcriptional initiation profiling(92) will be needed to resolve TSS usage across distinct astrocyte subtypes, and in vivo TSS mapping in models of acute and chronic inflammation will be necessary to capture the full complexity of astrocyte regulatory dynamics in the brain microenvironment.

In conclusion, this work provides a mechanistic foundation for understanding how astrocytes execute specialized inflammatory gene expression programs, contributing to their unique roles in brain function and disease.

Using csRNA-seq to map transcriptional initiation at single-base-pair resolution in astrocytes, we define transcribed regulatory elements and establish a TSS-based regulatory grammar in which key astrocyte TF motifs influence both transcription initiation efficiency and TSS selection. We demonstrate how lineage-determining TFs direct NF-κB genomic targeting to define cell type-specific inflammatory enhancer repertoires and establish that astrocyte-specific TSRs are selectively enriched for CNS disorder risk variants. Together, our findings reveal that the regulatory logic governing astrocyte-specific inflammatory responses is encoded at the level of transcription initiation itself.

A key result of this study is a spatial regulatory grammar at active TSSs in astrocytes, consistent with recent work demonstrating position-dependent TF function in human regulatory elements(3, 69). Our findings extend this model to the astrocyte inflammatory response. First, NF-κB, AP1, and IRF were preferentially positioned upstream of IL-1B-induced TSSs, indicating that the spatial arrangement of sequence-specific TFs is critical for efficient recruitment of RNAPII and initiation of transcription during neuroinflammation. Second, the internal architecture of TSS usage patterns is itself regulated in a TF-specific manner. Specifically, TEAD and NF-κB motifs were preferentially associated with stimulus-dependent shifts in TSS selection, supporting a model in which certain TFs determine transcriptional precision, directing where within a regulatory element the pre-initiation complex assembles(70–74). In contrast, NFI motifs were primarily associated with changes in overall transcriptional magnitude without corresponding shifts in TSS patterns, consistent with models in which distinct classes of TFs independently regulate RNAPII recruitment and productive elongation(75–77).

The comparison between astrocytes and macrophages revealed that despite substantial overlap in stimulus-induced mRNA transcripts, the underlying enhancer repertoires were predominantly divergent with only ∼25% of induced TSRs shared. This finding supports a mechanistic model by which universal inflammatory TFs such as NF-κB can mediate specialized regulatory programs in different cell types^83^. We show that lineage-determining factors establish cell type-specific chromatin landscapes that define the sites to which stimulus-responsive TFs are preferentially recruited. In astrocytes, NF-κB is recruited to sites co-bound by NFI and TEAD, whereas in BMDMs it is directed by PU.1 and C/EBP factors. This mechanism aligns with previous work demonstrating the combination of lineage-specific TFs prime cell type-specific landscapes in macrophages and B cells(38). Our ChIP-seq data showing stimulus-induced co-recruitment of NFIA and TED4 to NF-κB-enriched sites further corroborate this collaborative binding model.

These findings also identify TEAD4 as a previously uncharacterized regulator of astrocyte inflammatory gene expression. TEAD factors canonically function as transcriptional effectors of YAP/TAZ downstream of the Hippo pathway(79–81), and while YAP has been implicated in astrocyte differentiation, it generally attenuates rather than promotes neuroinflammation(81–86). Our findings raise the possibility that TEAD4 contributes to astrocyte inflammatory gene regulation through a mechanism that is at least partially independent of canonical YAP co-activator activity. One candidate mechanism is the TRAF6-YAP-TEAD4 signaling axis, previously described in other cellular contexts(87), which could provide a route for IL-1B to activate TEAD4 in astrocytes independently of Hippo pathway status and warrants further investigations.

Integrating our astrocyte TSR map with human GWAS data reveals that the regulatory code defined here is directly relevant to human disease genetics. Transcribed mouse astrocyte TSRs were approximately twice as likely to correspond to functionally validated enhancers in human CRISPRi screens_­_compared to non-transcribed accessible regions, establishing active transcription as a robust predictor of regulatory function across species. Leveraging this conservation, we show that astrocyte TSRs are significantly enriched for heritability of CNS disorders, including AD, bipolar disorders and neuroticism, while macrophage TSRs were enriched for variants associated with hematological and immune traits, a pattern that directly mirrors the known biological roles of each cell type. The specificity of these associations support the broader principle that disease-associated non-coding variants preferentially disrupt cell type-specific regulatory elements(88–91).

Several limitations of this study should be noted. The regulatory mechanisms identified here are based on cooccupancy and motif evidence, and direct perturbation experiments will be needed to establish causal roles for NFIA and TEAD4 in governing the astrocyte inflammatory response. Additionally, the csRNA-seq experiments were performed in bulk primary astrocyte cultures stimulated with a single cytokine (IL-1B), which may not fully recapitulate the diversity of astrocyte states and regulatory responses present in vivo, where astrocytes are exposed to a combination of signals from neurons, microglia and other cell types, and where epigenomic states may differ from those of cultured cells. Future studies incorporating single-cell transcriptional initiation profiling^97^ will be needed to resolve TSS usage across distinct astrocyte subtypes, and in vivo TSS mapping in models of acute and chronic inflammation will be necessary to capture the full complexity of astrocyte regulatory dynamics in the brain microenvironment.

In conclusion, this work provides a mechanistic foundation for understanding how astrocytes execute specialized inflammatory gene expression programs, contributing to their unique roles in brain function and disease.

## Data and code availability

Raw and processed csRNA and ChIP sequencing data generated in this study have been deposited in the Gene Expression Omnibus database under accession number GSE270639 and data analysis tutorials are available at http://homer.ucsd.edu/homer/ngs/csRNAseq/index.html. The code used for the GWAS enrichment analysis and LDSC is available at https://github.com/ftlabucsd/Astrocytes-TSR. Additional datasets include mouse BMDM ChIP-seq (GSE109965), mouse BMDM csRNA-seq (GSE135498), mouse BMDM ATAC-seq data (GSE119691), mouse astrocytes ATAC-seq and H3K27Ac ChIP-seq (GSE185215), mouse astrocytes/neurons methylation (GSE140271), human astrocytes TT-seq (GSE253099), human astrocytes ATAC-seq (GSE252965).

## Acknowledgements

We would like to acknowledge Havilah Taylor for technical assistance with mouse colony management. This work was funded by the National Institute of Health (5R21DA056177 to C.B. and F.T.; 1R21DA060455 to C.B., S.H. F.T; 5U01DA051972 to C.B and F.T.; R35GM149520 to C.B.)

## Author contributions

Conceptualization, FT, CB, AK; Investigation, AK, DS, NP; Formal Analysis, CB, YZ, NF, AK; Methodology, AK, SH, CG, FT, CB; Writing-Original Draft, FT, CB; Writing-Review & Editing, FT, CB, AK, YZ, NF, DS, NP, CG, SH; Supervision, FT, CB.

## Materials and Methods

### Antibodies

Antibodies used in immunohistochemistry are anti-GFAP (Millipore Cat# MAB360, RRID: AB_11212597) 1:500 and Donkey anti-Mouse (Thermo Fisher Scientific Cat# A-21203, RRID: AB_141633) 1:1000. Antibodies used in ChIP-seq are anti-Pan-TEAD (Cell Signaling, #13295, RRID:AB 2687902), anti-NFIA (Millipore-Sigma, HPA006111) and RNAPII (Cell Signaling, #14958).

### Primary Astrocyte Cultures

Primary astrocyte cultures were established by dissecting cortical tissues from P1 C57BL/6J mouse pups (RRID: IMSR_JAX:000664). After tissue homogenization, the isolated astrocytes were then plated at a density of 130,000 cells/cm2 and maintained for 19 days in vitro (DIV) to allow astrocytes to mature. DMEM-F12 media with heat-inactivated 10% FBS was changed every 3 days. At DIV 7, cultures were split after shaking overlaying microglia, if present. Two days before collection, the astrocytes were plated at a density of 500,000 cells per well on 4-well chamber slides to perform immunocytochemistry or a density of 5.5 million cells per 15 cm dish to be treated with 10ng/mL of recombinant human interleukin 1 beta (IL-1B) (Invivogen, Cat#rcyec-h) for 1hr before sample collection. A step-by-step protocol was previously published(93).

### Immunocytochemistry

The cells were plated on 4-well chamber slides (Thermo Fisher Scientific Cat#PEZGS0416) at 500,000 cells per well and fixed with 4% formaldehyde (Thermo Fisher Scientific Cat#28906) at RT for 15 min. After permeabilization with 0.3% Triton X-100 for 5 min at RT, blocking in super blocking buffer (Thermo Fisher Scientific Cat#37515) was performed for 1 hr at RT. The primary antibodies were diluted in Ab Dilution Buffer (10% Superblock + 0.1% Tween in PBS 1X) and then incubated overnight at 4C. The next day, the cells were blocked again for one hour at RT. Secondary Antibodies were diluted 1:1000 in Ab Dilution Buffer and incubated for one hour at RT. The nuclei were stained with Hoechst 33342 (Thermo Fisher Scientific Cat#H3570) 1:1000 in PBS for 5 min at RT. The slides were then mounted with Prolong Glass antifade Mountant (Invitrogen Cat#P36984) and left to dry for 24 hr before imaging at the Keyence digital fluorescent microscope (Keyence Cat#BZX800). ImageJ (RRID:SCR_003070) software was used to count the cells that showed the signal. A step-by-step protocol was previously published(94).

### Total RNA extraction and cDNA synthesis for RT-qPCR

RNA was extracted using TRIzol (Thermo Fisher Scientific Cat#15596018) following the manufacturer’s instructions. 100 ng of RNA was digested with DNAse I (Thermo Fisher Scientific Cat#K1681). RNA was then reverse transcribed to cDNA by Maxima H Minus First Strand cDNA Synthesis Kit with dsDNase (Thermo Fisher Scientific Cat#K1681). Real-time quantitative polymerase chain reaction was done using SYBR® Green PCR Master Mix (Thermo Fisher Scientific Cat#4309155) and carried out in Bio-Rad CFX384 Real-Time Detection System (RRID:SCR_018057) and analyzed with CFX Maestro software (RRID:SCR_018057). The full primer sequences are the following: *Csf (*Forward-GCCTTGGAAGCATGTAGAGG, Reverse-CTGGTAGTAGCTGGCTGTCA*), Ccl20 (*Forward-TGTACGA-GAGGCAACAGTCG*, Reverse-TCTGCTCTTCCTTGCTTTGG), Cxcl1* (Froward-CCTATCGCCAATGAGCTGC, Reverse-CTCGCGACCATTCTTGAGTG), *Irf1* (Forward-ACAGACGAGGATGAGGAAGG, Reverse-GGCTGTCAATCTCTGGTTCC)*, Tnfaip3* (Forward-GCTCAACTGGTGTCGTGAAG, Reverse-GTTCCGAG-TGTCTGTCTCCT)*, Tnfa* (Forward-CCACGCTCTTCTGTCTACTGAACT, Reverse-AGGGTCTGGGCCATAGAACT)*, Il6* (Forward-ACCAGAGGAAATTTTCAATAGGC, Reverse-TGATGCACTTGCAGAAAACA). RT-qPCR values were expressed as means +/- standard error of the mean (SEM), and significant results were reported with a p<0.05. Data visualization and statistical analysis were performed using GraphPad Prism 9 (RRID:SCR_002798). An unpaired t-test was used to compare DMSO vs. IL-1B in RT-qPCR experiments for each target gene tested.

### RNA-seq library preparation for sequencing

RNA-seq libraries were generated from total RNA that underwent an in-column DNase I treatment for 30 minutes using the RNA Clean & Concentrator-5 kit (Zymo Cat#R1013). 1ug of DNase-treated RNA was used to generate libraries following the manufacturer’s protocol for the NEBNext® Ultra™ II Directional RNA Library Prep Kit for Illumina (NEB Cat#E7760S) with the Poly(A) mRNA Magnetic Isolation Module (NEB Cat#E7490). After poly(A) enrichment and fragmentation, RNA was synthesized into cDNA with NEBNext Adaptor diluted 1:20. NEB barcodes (NEB Cat#E6440), and the libraries were amplified by PCR for 8 cycles and sequenced on the Illumina NextSeq 500 (RRID:SCR_014983).

### Capped small RNA-seq library preparation for sequencing

csRNA-seq libraries were generated from ∼3μg of total RNA. First, the short RNAs of ∼20–60 nt were isolated after excision from a denaturing Novex TBE-Urea 15% Gel (Invitrogen Cat#EC68852BOX). An input sample corresponding to 16% of the volume of the purified short RNAs was saved for each sample. The 5’-capped RNAs containing a 3’-OH were enriched through a series of enzymatic reactions. Monophosphorylated RNAs were degraded by Terminator 5′-phosphate-dependent exonuclease (Lucigen Cat#TER51020). Next, 5′dephosphorylation by CIP (NEB Cat#M0525L) was followed by RNA 5′ Pyrophosphohydrolase (RppH) to convert the 5′ end of triphosphorylated RNA to a 5′ monophosphate RNA (NEB Cat#M0356S). Adapters were ligated to both the 5’ cap end and the 3’ end. The RNA was synthesized into cDNA with NEB barcodes (NEB Cat#E7500S), and the libraries were amplified by PCR for 14 cycles and sequenced on the Illumina NextSeq 500 (RRID:SCR_014983).

### Chromatin Immunoprecipitation sequencing

Chromatin immunoprecipitation sequencing (ChIP-seq) was performed on primary astrocytes using a double-crosslinking protocol. Cells were sequentially fixed with freshly prepared 2 mM disuccinimidyl glutarate (DSG) in 1× PBS followed by formaldehyde, then quenched with 125 mM glycine and washed with cold PBS. Nuclei were isolated in a hypotonic lysis buffer with a 30 min incubation. Chromatin was sheared using a Pixul sonicator (Active Motif) in RIPA containing 0.4% SDS. Antibodies were pre-bound to Dynabeads Protein Protein G (Thermo Fisher Scientific) and incubated with sheared chromatin overnight. Immunoprecipitates were washed and bound chromatin was eluted, treated with RNase A and proteinase K, and crosslinking was reversed at 65°C. DNA was purified using PEG/NaCl bead-based size selection and eluted in Tris-Tween buffer. Input DNA was processed in parallel from aliquots of sonicated chromatin. Sequencing libraries were prepared using NEBNext Ultra II DNA Library Prep Kit for Illumina (E7645S). Libraries were PCR-amplified using indexed primers, purified by bead cleanup, and assessed for size distribution by TapeStation. Library concentration was measured using a high-sensitivity Qubit assay prior to sequencing on the Illumina NextSeq 500.

### Bioinformatic analysis of NGS data

TSS location and activity levels were determined using csRNA-seq and generally analysed using HOMER2(3, 38). Additional information, including analysis tutorials are available at http://homer.ucsd.edu/homer/ngs/csRNAseq/index.html. Other data types, including RNA-seq, ChIP-seq, ATAC-seq, and DNA methylation were also largely processed with HOMER, and are also described below.

csRNA-seq (small capped RNAs, ∼20–60 nucleotides) and total small RNA-seq input sequencing reads were trimmed of their adapter sequences using HOMER (‘homerTools trim −3 AGATCGGAAGAGCACACGTCT -mis 2 -minMatchLength 4 -min 20’) and aligned to the mouse GRCm38/mm10 genome using STAR (v.2.7.10a)(95) with the default parameters. Only reads with a single, unique alignment (MAPQ ≥ 10) were considered in the downstream analysis.

Furthermore, reads with spliced or soft clipped alignments were discarded to ensure accurate TSSs. Transcription Start site Regions, or TSRs, representing loci with significant transcription initiation activity from one or more individual TSSs on the same strand from the same regulatory element (that is, peaks in csRNA-seq), were defined using HOMER’s findcsRNATSR.pl tool, which uses short input RNA-seq, traditional total RNA-seq and annotated gene locations to find regions of highly active TSSs and then eliminate loci with csRNA-seq signals arising from non-initiating, high-abundance RNAs that nonetheless are captured and sequenced by the method (further details are available in a previous study(6)). Replicate experiments were first pooled to form meta-experiments for each condition (e.g. NT, IL-1B) before identifying TSRs. Annotation information, including gene assignments, promoter distal, stable transcript and bidirectional annotations are provided by findcsRNATSS.pl. To identify differentially regulated TSRs, TSRs identified in each condition were first pooled (union) to identify a combined set of TSRs represented in the dataset using HOMER’s mergePeaks tool using the option ‘-strand’. The resulting combined TSRs were then quantified across all individual replicate samples by counting the 5′ ends of reads aligned at each TSR on the correct strand. The raw read count table was then analysed using DESeq2(96) to calculate normalized rlog-transformed activity levels and identify differentially regulated TSRs (defined as adj. P-value < 0.05, Log2 fold-change > 1). Changes in TSS initiation patterns (or ‘shape’) were calculated using HOMER’s compareTSS.pl, which implements a modified version of the WIP score to assess differences in TSS selection between two samples(54). Briefly, the WIP is calculated by first normalizing the TSS levels in a TSR to a total of 1 in each of the conditions being compared. Then, for each initiation position, the difference is calculated as the absolute value of the difference between normalized signal levels at each position from [-25,+25] linearly scaled to the distance from the initiation site, penalizing differences that occur further from the reference TSS. WIP scores are also calculated between replicate experiments to create a NULL distribution, which is then used to assess the significance values for the comparison.

RNA-seq data was processed similarly to that described above for csRNA-seq, except that read counts per gene were calculated using GENCODE defined transcripts using HOMER’s analyzeRepeats.pl quantification tool, and differentially expressed genes were calculated using DESeq2(96). Metascape was used to calculate pathway enrichment scores for differentially regulated genes(29). DNA methylation calls were taken directly from supplementary files supplied to NCBI GEO (GSE140271) and average cytosine methylation calls around loci were found using HOMER’s annotatePeaks.pl program. ChIP–seq and ATAC-seq reads generated by this study and from previously published studies were aligned to the mouse genome (GRCm38/mm10) using STAR with the default parameters. Only reads with a single, unique alignment (MAPQ ≥ 10) were considered in the downstream analysis. ChIP–seq peaks were determined using HOMER’s findPeaks program using ‘-style factor’, while ATAC-seq peaks were defined using ‘-style atac’. Quantification of ChIP–seq reads associated with peaks, annotation to the nearest annotated gene TSS, calculation of TF binding site presence (−100 to +100 relative to the peak center) and visualization of normalized read pileups for the genome browser were all conducted using HOMER.

In all cases, normalized genome browser visualization tracks for RNA-seq, csRNA-seq, and ChIP/ATAC-seq data were generated using HOMER’s makeUCSCfile program using the ‘-rna’, ‘-tss’, or ‘-chip’ options, respectively, and visualized using the UCSC Genome Browser(97). Annotation of TSS/TSR locations to the nearest gene was performed using HOMER’s annotatePeaks.pl program using REFSEQ as the reference annotation. Histograms and quantification of csRNA, ATAC, ChIP, or other sequencing reads were performed using HOMER’s annotatePeaks.pl program. Homologous (i.e. conserved) TSRs were identified using the UCSC Genome Browser’s liftover tool (mm10toHg38) and required at least 10% of the mapped element to be present in the human genome.

Motif enrichment calculations were performed using HOMER, which scores motif enrichment by comparing the frequency of motif instances against a random set of genomic background sequences that are chosen to match the GC-properties of the target sequences. TSRs are analyzed from −150 to +50 relative to the primary TSS, while ChIP/ATAC-seq peak regions are analyzed from −50 to +50 relative from the peak center position. Predicted motif instances and densities were calculated using HOMER’s annotatePeaks.pl tool, and enrichment heatmaps were created using Cluster 3.0/Java TreeView(98). GREAT(47) version 4.0.4 was used to identify pathways enriched in TSRs. TSRs lifted over from mm10 to hg19 were used as inputs of GREAT, with the background being the whole genome. The following association rules were applied: basal+extension: 5000 bp upstream, 1000 bp downstream, 1000000 bp max extension; curated regulatory domains included. Pathways with hypergeometric test FDR < 5% were considered significant.

#### GWAS enrichment analysis

We downloaded the NHGRI-EBI GWAS Catalog (v1.0.2, March 28, 2024) from https://www.ebi.ac.uk/gwas/docs/file-downloads. GWAS associations were processed to define unique loci by combining chromosome and base-pair position (CHR_ID:CHR_POS). Trait-study combinations were defined as unique pairs of a reported trait and PubMed ID (TRAIT.PUBMEDID). Entries lacking genomic coordinates were removed. To reduce redundancy due to linkage, SNPs within 5 kb of a preceding SNP within the same trait-study combination and chromosome were excluded after sorting by genomic position. We then retained only SNPs meeting nominal significance (P ≤ 0.05) and filtered trait-study combinations to include those with at least 10 significant SNPs. Trait-study combinations with a total sample size ≤20,000 were excluded. After filtering, 2,025 trait-study combinations (from 1,304 traits and 750 publications) were retained for downstream analysis. For enrichment analysis, we tested whether GWAS loci overlapped TSRs identified by csRNA-seq. Overlap between GWAS loci and TSRs was computed for each trait-study combination. Enrichment was assessed using a hypergeometric test, where: the number of successes corresponds to overlapping loci, the background universe consists of all unique GWAS loci across filtered datasets and TSR loci, and the sample size corresponds to the number of loci associated with each trait-study combination. FDR values were calculated using the Benjamini-Hochberg procedure. Statistical significance was defined as FDR < 0.05.

#### Linkage disequilibrium score regression (LDSC)

To assess enrichment of human GWAS signal within TSRs, we performed stratified LDSC. TSRs identified from mouse astrocytes and macrophages were first converted from mm10 to hg38 genomic coordinates using the UCSC liftOver tool on the UCSC Genome Browser, requiring a minimum remapping fraction of 0.5(99). Lifted TSR regions (including centered 100 bp windows) were used as foreground annotations. As background, we incorporated DNase I hypersensitivity regions from 53 epigenomes (ENCODE Honeybadger2 release (https://personal.broadinstitute.org/meuleman/reg2map/HoneyBadger2_release/). Annotation files were generated using *make_annot.py* from the LDSC software package, intersecting TSR BED files with SNP coordinates from the 1000 Genomes Project Phase 3 European reference panel (PLINK format, chromosomes 1-22)(100). LD scores were computed for each annotation using *ldsc.py* --l2, with a 1 cM LD window and restriction to HapMap3 SNPs(101). Partitioned heritability analyses were then performed using l*dsc.py --h2-cts*, incorporating baseline LD annotations (baseline model) and regression weights derived from HapMap3 SNPs. Custom TSR annotations were included as cell-type-specific annotations via the *--ref-ld-chr-cts* framework.

GWAS summary statistics included 297 traits from Zhang et al. (2021)(61) and an additional 35 neuropsychiatric and autoimmune traits. Summary statistics were harmonized and converted into LDSC-compatible format using *munge_sumstats.py*.

For each trait, we estimated the enrichment coefficient and corresponding P value for astrocyte and macrophage TSR annotations relative to the baseline model. Multiple testing correction was performed using the Benjamini–Hochberg procedure across all traits, and annotations with false discovery rate (FDR) < 0.1 were considered significant.

**Figure S1:**
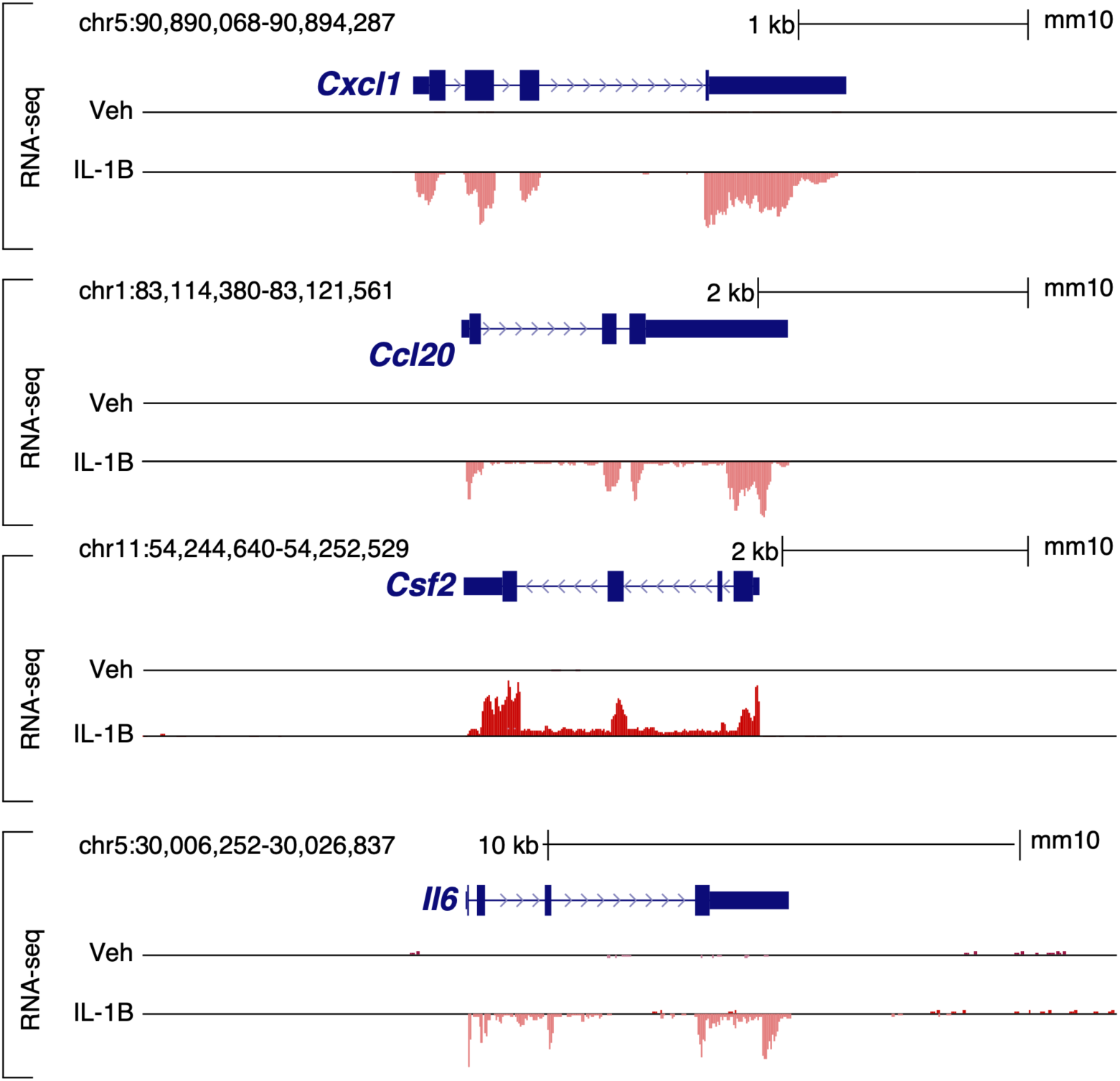
Representative RNA-seq browser tracks of IL-1B-responsive genes in primary astrocytes. Genome browser views showing RNA-seq signal at the *Cxcl1, Ccl20, Csf2,* and *Il6* loci in astrocytes treated with vehicle or IL-1B. These representative examples illustrate the robust transcriptional induction of canonical inflammatory genes following IL-1B stimulation. Gene models and genomic coordinates are shown in *mm10*.

**Figure S2:**
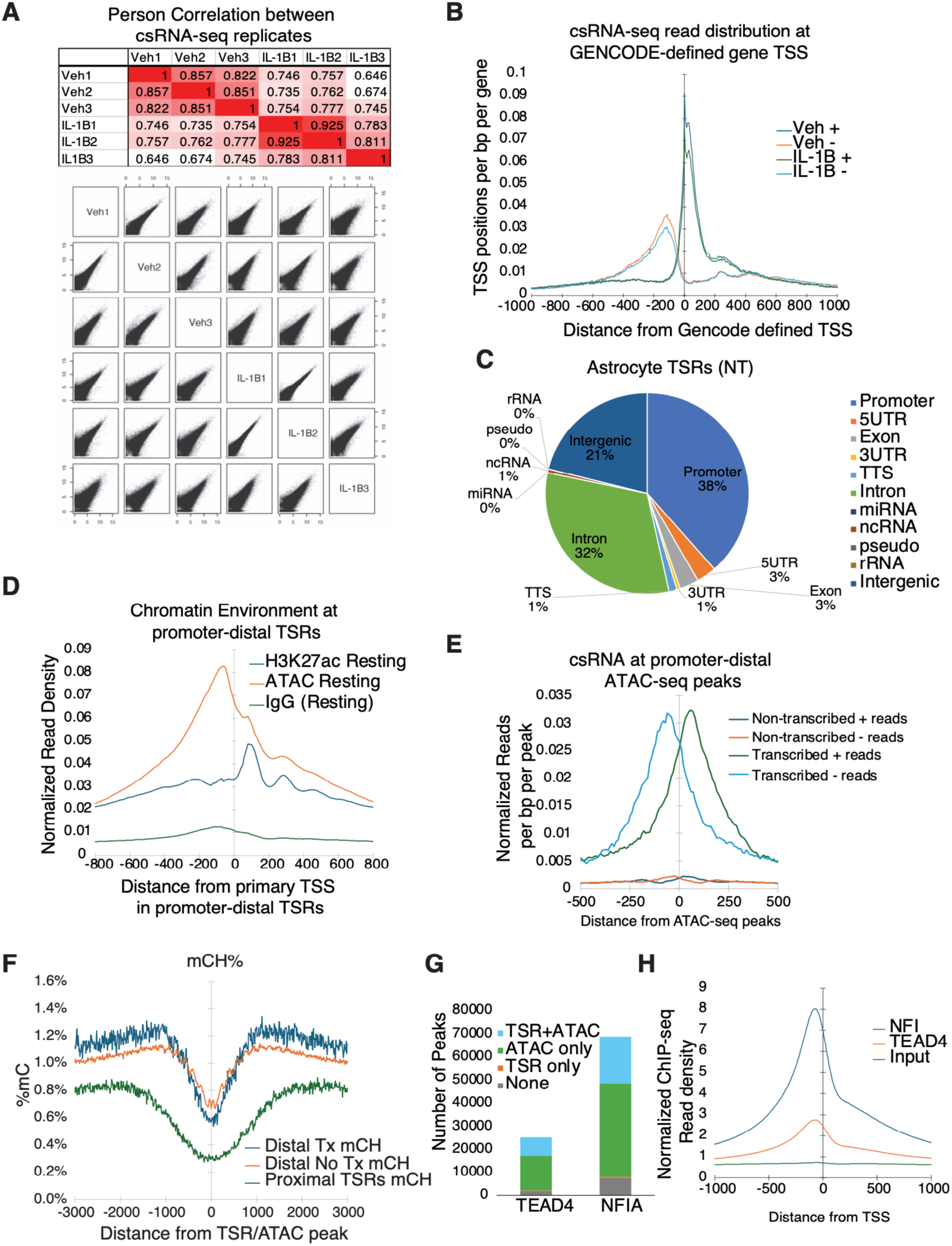
Features of astrocyte TSRs and associated chromatin environment. (A) Pairwise correlation analysis of csRNA-seq replicates from untreated and IL-1B-treated astrocytes. (B) Distribution of strand-specific csRNA-seq reads around GENCODE-annotated TSSs, showing strong enrichment at annotated transcriptional start sites. (C) Genomic annotation of astrocytes TSRs, partitioned across promoter, intronic, intergenic, and other genomic features. (D) Average chromatin profiles at promoter-distal TSRs, showing ATAC-seq and H3K27ac enrichment around transcribed regulatory elements. (E) Average csRNA-seq signal centered on promoter-distal ATAC-seq peaks, comparing transcribed and non-transcribed accessible regions (No Tx: 225,150 peaks, Tx: 14,836 peaks). (F) Average mCH profiles at promoter-proximal TSRs, promoter-distal transcribed TSRs, and non-transcribed distal accessible regions in astrocytes (data from^50^). (G) Distribution of NFIA and TEAD4 ChIP-seq peaks found overlapping TSRs and ATAC-seq peaks in astrocytes. (H) ChIP-seq read density for NFIA and TEAD4 centered on csRNA-seq defined TSRs, showing transcription factor binding immediately upstream of the primary TSS.

**Figure S3:**
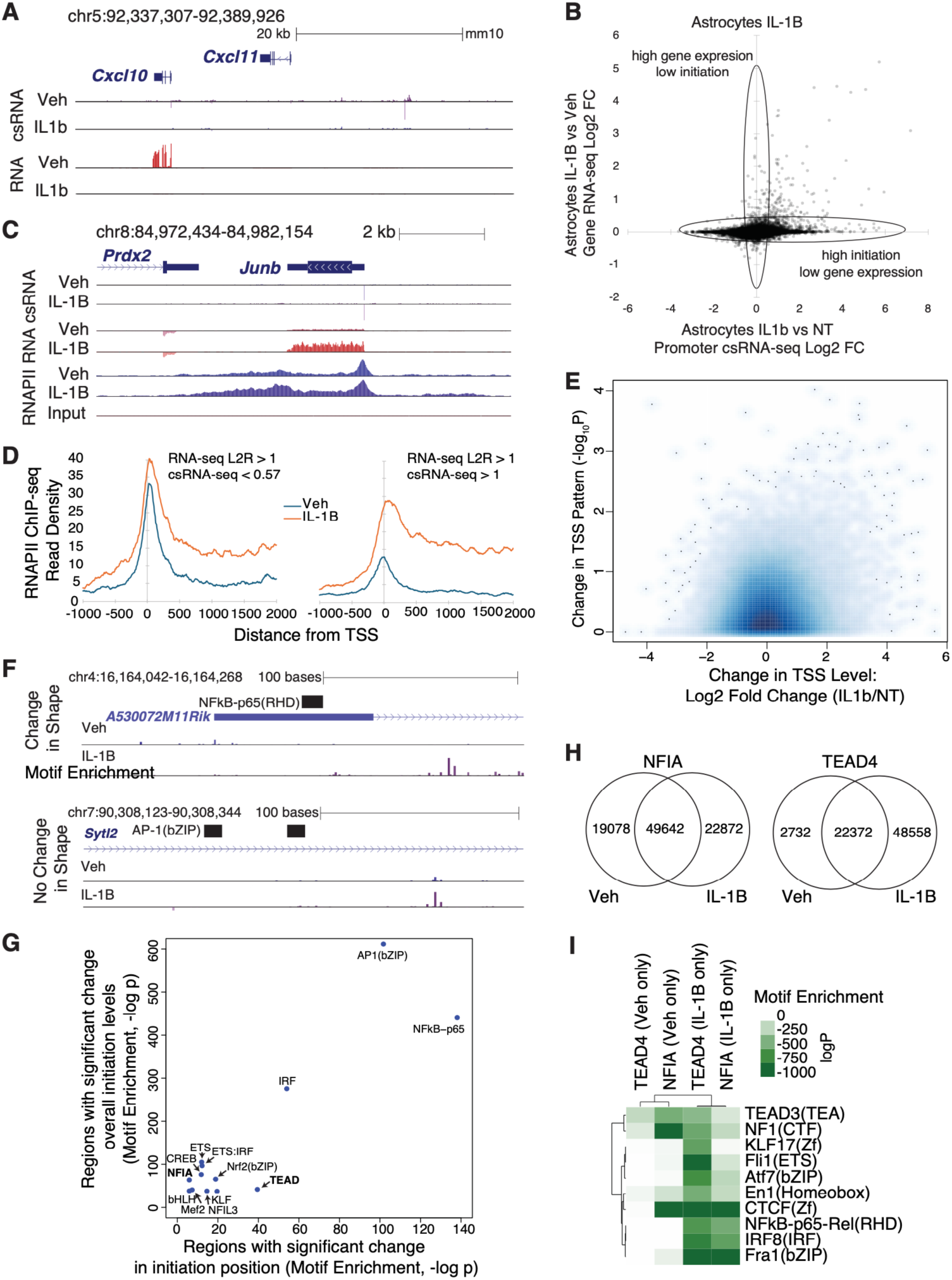
Distinct effects of IL-1B on transcription initiation level and TSS architecture. (A) Genome browser example of an IL-1B-induced locus (Cxcl10) in astrocytes, showing increased transcription initiation after stimulation. (B) Scatter plot comparing IL-1B-induced Log2 csRNA-seq changes at the promoter vs. RNA-seq changes across genes, highlighting genes regulated primarily at initiation (along x-axis) versus those showing stronger changes at the mRNA level (along y-axis). (C) Genome browser tracks at the Junb locus showing increased gene expression and RNAPII elongation in the gene body with limited change in promoter initiation and RNAPII promoter levels, consistent with regulation being mediated primarily at the level of transcription elongation rather than increased initiation. (D) RNAPII ChIP-seq levels at the promoters of IL-1B induced genes stratified by genes with minimal versus strong increases in csRNA-seq initiation activity. (E) Scatter plot comparing changes in overall TSR levels (Log2 Fold change, NT vs. IL-1B) versus their WIP score significance (the −Log10 p-value), identifying TSRs with altered initiation patterns independent of changes in total transcriptional output. (F) Representative examples of TSRs exhibiting a strong change in initiation pattern (top, WIP score −1.61, Lg10 p-value = 9.54e-05) versus a strong change in overall initiation levels with minimal change in initiation shape(WIP score −0.13, Log10 p-value = 0.84). (G) Scatter plot of TF motif enrichment in TSRs with significant changes in overall activity versus changes in TSS positions, highlighting differential associations of NF-κB, TEAD, and NFI motifs with these TSRs classes. (H) Venn diagram showing the overlap of ChIP-seq peaks for NFIA and TEAD4 before and after IL-1B stimulation in astrocytes. (I) Motif enrichment analysis of condition-specific (either Veh or IL-1B) TEAD4- and NFIA-bound regions, showing IL-1B-specific enrichment for inflammatory TF motifs, including NF-κB, IRF (IRF8), and AP1 (i.e. Fra1).

**Figure S4:**
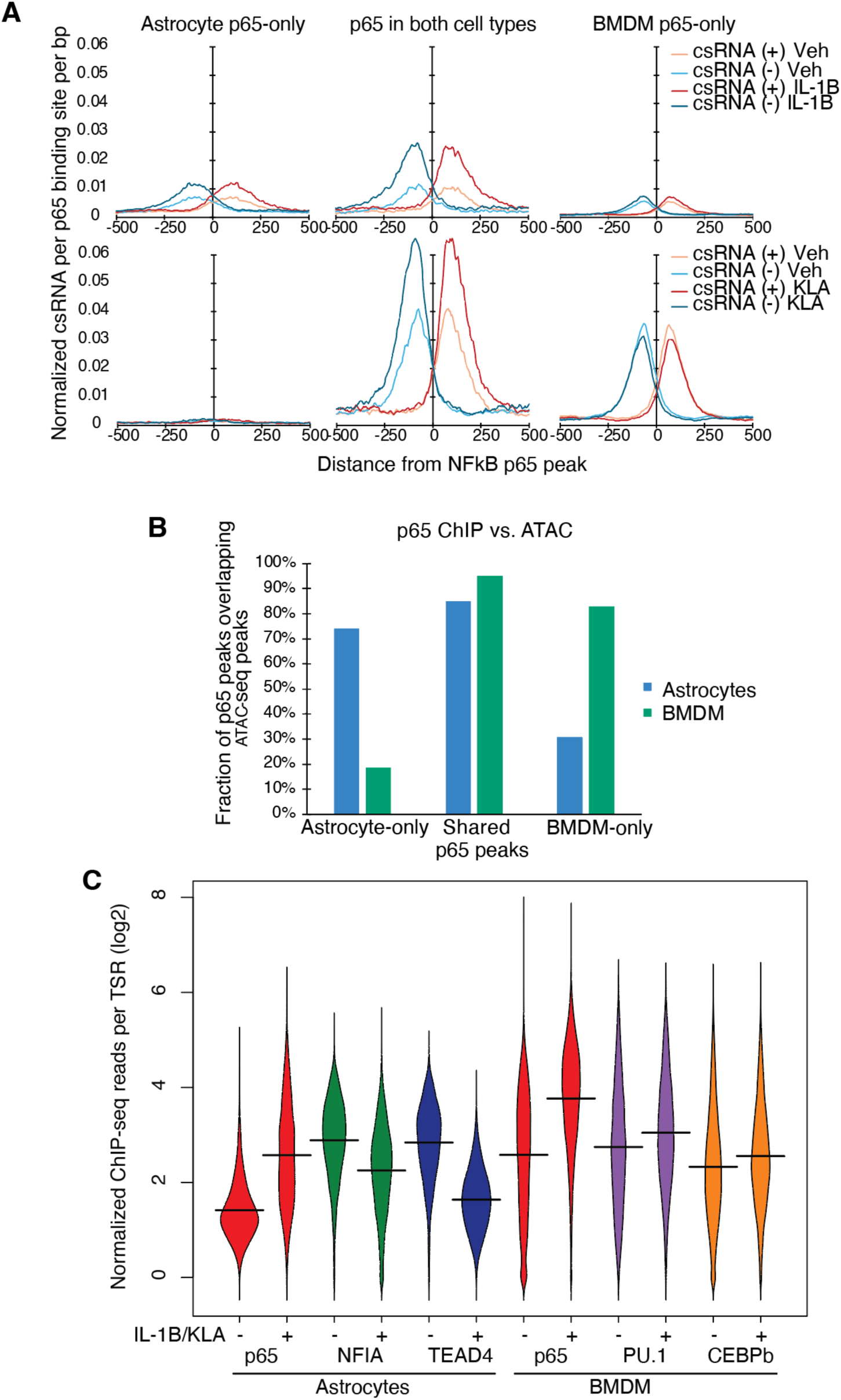
NF-κB binding is cell-type restricted and linked to pre-existing chromatin and lineage TF occupancy. (A) Average eRNA signal centered on astrocyte-specific, shared, and BMDM-specific NF-κBp65 peaks, showing cell type-matched induction of regulatory transcription at p65 bound sites. (B) Fraction of astrocyte-specific, shared, and BMDM-specific p65 peaks overlapping accessible chromatin regions in astrocytes or BMDMs, indicating that NF-κB recruitment occurs preferentially at cell-type-specific open chromatin regions. (C) Violin plots showing increase in ChIP-seq signal for p65, NFIA, TEAD4 at astrocyte IL-1B-induced TSRs and for p65, PU.1, and CEBPβ in macrophage KLA-induced TSRs.

**Figure S5:**
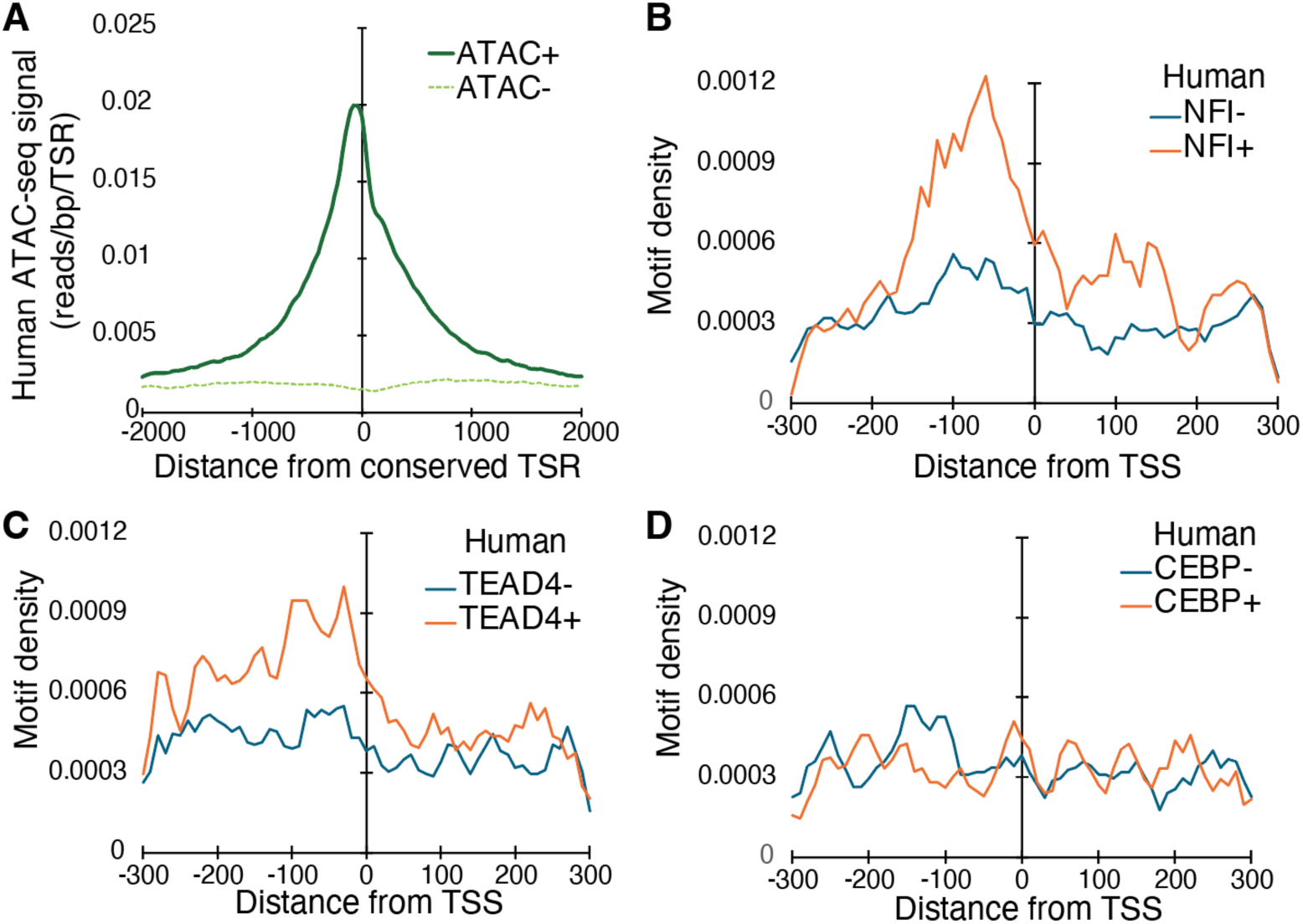
Human accessibility and motif architecture at conserved astrocyte TSRs. (A) Distribution of ATAC-seq reads in human astrocytes at conserved mouse TSRs that overlap human ATAC-seq peaks (ATAC+) versus those that do not (ATAC-). (B) Average human NFI motifs centered on conserved mouse TSRs, stratified by whether the homologous human region is accessible or not. (C) Average human TEAD4 motifs centered on conserved mouse TSRs, stratified by whether the homologous human region is accessible or not. (D) Average human CEBP motifs centered on conserved mouse TSRs, stratified by whether the homologous human region is accessible or not.

**Figure S6:**
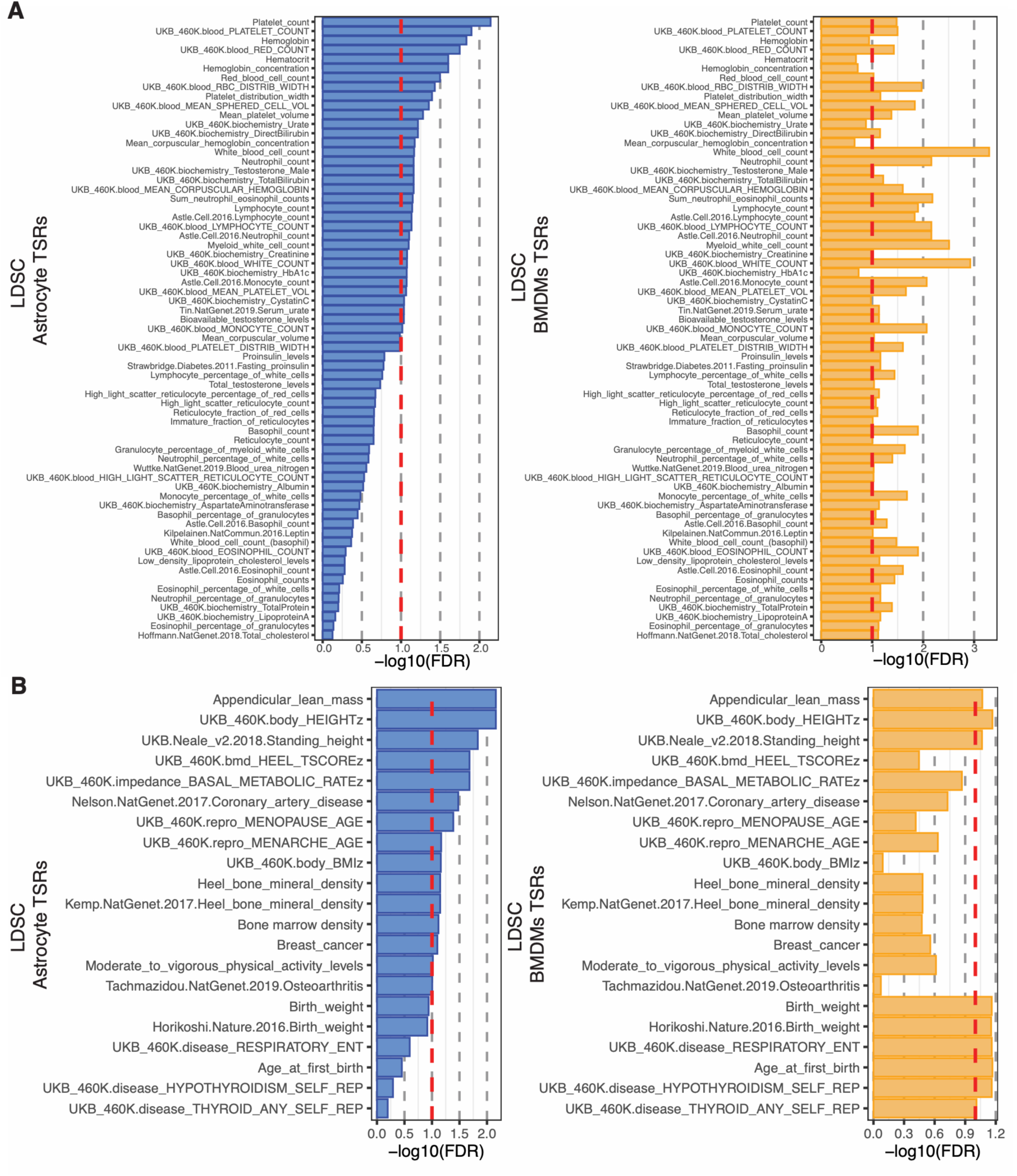
LDSC-based enrichment of additional GWAS traits. (A) LDSC regression enrichment for hematological traits across conserved astrocyte TSRs (left) or BMDMs TSRs (right). (B) LDSC regression enrichment for remaining traits across conserved astrocyte TSRs (left) or BMDMs TSRs (right).

## Supplemental Information

**Supplemental Table 1: Dataset Statistics**

List of experiments and read counts for data generated in this study.

**Supplemental Table 2: RNA-seq DEGs**

Two worksheets describing genes induced or repressed by IL-1B in primary mouse astrocytes. The final two columns describe genes that are also significantly regulated by KLA in BMDMs.

**Supplemental Table 3: csRNA-seq TSRs**

Table describing properties of the 69,435 TSRs identified in mouse astrocytes. Each TSR entry contains information about its position in the mm10 genome, annotation to nearby genomic features and genes, csRNA-seq normalized count information and differential regulation by IL-1B, overlap annotation with ATAC-seq, NF-κB p65 binding, and BMDM KLA regulation, changes in TSS initiation site selection (WIP2 scores), and human conservation information.

**Supplemental Table 4: GWAS enrichment**

GWAS enrichment results for overlap between csRNA-seq-defined TSRs and filtered GWAS Catalog loci. Results are reported for each trait-study combination, including total loci, overlapping loci, enrichment statistics, empirical P values, and FDR-adjusted significance. The first sheet includes all tested trait-study combinations, and the second sheet reports the subset of traits related to neurological and neuropsychiatric phenotypes.

**Supplemental Table 5: LDSC**

The table reports LDSC enrichment statistics, regression coefficients, standard errors, nominal P values, and multiple-testing-adjusted significance values for each trait and TSR set (astrocytes, BMDMs).

